# Expansion and Comprehensive Evaluation of the Peacock Eye Disease Management (PedMan) System across Olive Cultivars Differing in Susceptibility to *Venturia oleaginea*

**DOI:** 10.64898/2026.11.08.765137

**Authors:** Yotam Gilat, David Ygzao, Dani Shtienberg, David Ezra

## Abstract

Peacock eye disease, caused by *Venturia oleaginea*, is an important foliar disease of olive that is primarily managed through fungicide applications. Integrating host resistance into disease-warning systems could improve spray timing and reduce unnecessary fungicide use. PedMan (Peacock eye disease Manager) was previously developed and validated for highly susceptible olive cultivars. The objectives of the present study were to expand PedMan to cultivars differing in susceptibility to *V. oleaginea* and to evaluate the expanded system under commercial grove conditions. A sensitivity analysis of the rainfall and temperature criteria used for highly susceptible cultivars was conducted to establish criteria for moderately susceptible and moderately resistant cultivars. The rainfall threshold of ≥15 mm was retained, whereas the temperature range triggering an infection alert was narrowed from 12–18 °C for highly susceptible cultivars to 13–17 °C for moderately susceptible and 14–16 °C for moderately resistant cultivars. The expanded system was subsequently evaluated in 33 field trials and observations under natural infection conditions. PedMan recommendations were classified as correct or incorrect based on statistical comparisons of leaf abscission between fungicide-treated and untreated trees. Overall, 30 of 33 recommendations were correct, resulting in an accuracy of 90.9%. Sensitivity was 87.0%, while specificity and positive predictive value were both 100%. These results demonstrate that incorporating cultivar susceptibility into PedMan broadens its applicability and provides reliable cultivar-specific fungicide recommendations. The system has potential to reduce unnecessary fungicide applications while maintaining effective management of peacock eye disease across olive cultivars differing in susceptibility.

## 1. Introduction

Peacock eye disease, also known as olive leaf spot, is one of the most important foliar diseases of olive (*Olea europaea*) and is caused by the fungus *Venturia oleaginea* (syn. *Spilocaea oleagina*). The disease is widely distributed throughout the Mediterranean Basin and other olive-growing regions and can cause substantial reductions in tree productivity (Graniti, 1993; Buonaurio et al., 2023). The disease primarily affects leaves, where characteristic circular, dark lesions surrounded by yellow halos develop on the upper leaf surface. Severe and repeated infections result in premature leaf abscission and extensive defoliation, reducing the photosynthetic capacity and vigor of affected trees and potentially leading to reduced flowering and yield (Graniti, 1993; Buonaurio et al., 2023). Disease development is strongly influenced by environmental conditions, particularly temperature, rainfall, and leaf wetness, which determine conidial release, dispersal, germination, and penetration into the leaf tissue (Graniti, 1993; Viruega and Trapero, 1999; Viruega et al., 2011; Obanor et al., 2011). Current disease management practices rely primarily on timely fungicide applications and the use of cultivars with reduced susceptibility to the pathogen. Copper-based fungicides remain the principal means of chemical control and are widely used as protectant treatments, while other fungicide groups, including dodine, strobilurins, and triazoles, have also demonstrated efficacy against the disease (Obanor et al., 2008; Buonaurio et al., 2023). However, effective chemical control depends strongly on application timing relative to infection events, and conventional calendar-based spray programs may result in unnecessary applications when environmental conditions are unfavorable for infection. The use of genetically resistant or less susceptible cultivars therefore represents an important component of integrated disease management and can reduce dependence on fungicides (López-Doncel et al., 1999; Buonaurio et al., 2023). Thus, combining cultivar resistance with environmentally based, risk-oriented fungicide applications provides an opportunity to improve disease control while reducing unnecessary fungicide use.

Polygenic or quantitative resistance to plant diseases is generally expressed as a continuous range of phenotypic responses rather than as a simple resistant–susceptible dichotomy. Such resistance is often referred to as partial, quantitative, horizontal, or non-race-specific resistance and is typically controlled by several genes, each contributing a portion of the overall resistance (Parlevliet, 1976; Agrios, 2005; Niks et al., 2015). For practical implementations, genotypes exhibiting quantitative variation in disease severity are often classified into discrete response classes, commonly including highly resistant (HR), moderately resistant (MR), moderately susceptible (MS), and highly susceptible (HS). This classification should be regarded as an operational categorization of a continuous resistance phenotype rather than as evidence for four genetically distinct resistance groups.

The response of *O. europaea* cultivars to *V. oleaginea* is quantitative and ranges from highly susceptible to highly resistant (Graniti, 1993; Buonaurio et al., 2023). To the best of our knowledge, no olive cultivar has yet been reported to be completely immune to the pathogen. Numerous studies conducted in different countries have evaluated the responses of local and international olive cultivars to the pathogen, including MacDonald et al. (2000) in New Zealand; López-Doncel et al. (1999), Moral et al. (2015) and Estudillo et al. (2025) in Spain; Ciccarese et al. (2002) in Italy; Mahfoud et al. (2018) in Syria; Abuamsha et al. (2013) in Palestine; Afiad-Lachger et al. (2020) and Najafi et al. (2023) in Iran; and Bernaschina et al. (2019) in Uruguay. In most cases, the classification of a given cultivar is consistent across studies. For example, cv. Souri was classified as highly susceptible in all studies in which it was evaluated (MacDonald et al., 2000; Mahfoud et al., 2018; Najafi et al., 2023; Ygzao et al., 2026a; 2026b)). However, it should be noted that, in some cases, the response of a given cultivar to the pathogen varies among studies and geographical regions. For example, in New Zealand and Uruguay, cv. Barnea was classified as highly or moderately susceptible (MacDonald et al., 2000; Obanor et al., 2011; Bernaschina et al., 2019), whereas in Spain it was categorized as “doubtful” (López-Doncel et al., 1999). In Israel, cv. Barnea has been classified as moderately resistant. It should also be emphasized that, for a given cultivar, disease severity may vary considerably depending on local environmental conditions and inoculum pressure. Thus, a highly susceptible cultivar may exhibit only slight infection under conditions unfavorable for pathogen development or when the initial inoculum level is low. Under such conditions, highly resistant cultivars may remain completely uninfected. Conversely, under environmental conditions highly conducive to pathogen development and in the presence of abundant inoculum, even highly resistant cultivars may develop moderate or, in some cases, severe disease.

Our recent studies have provided new insights into the epidemiology and management of peacock eye disease under Israeli conditions. We identified two distinct infection periods during the growing season, occurring in autumn, early winter and in late spring, early summer, and demonstrated that *V. oleaginea* is effectively monocyclic within a growing season, completing one major disease cycle per season (Ygzao et al., 2026c). Field experiments conducted in commercial olive groves further showed that one or a few fungicide applications in the autumn were sufficient to effectively suppress disease development during the same season (Ygzao et al., 2026b), supporting the epidemiological findings and indicating that fungicide applications can be timed according to infection risk rather than fixed calendar intervals. Based on these findings and our studies of the environmental conditions governing infection (Ygzao et al., 2026a), we developed PedMan (Peacock eye disease Manager), a decision support system (DSS) designed to identify infection events and provide recommendations for optimal fungicide application timing. PedMan integrates rainfall and temperature data to predict infection events, defined as periods favorable for conidial liberation, dispersal, germination, and penetration. For highly susceptible cultivars, PedMan generates an infection alert when rainfall reaches ≥15 mm in a single day or accumulates to ≥15 mm over consecutive days, provided that the average minimum temperature during the rainy days is 12–18 °C (Ygzao et al., 2026a; Ezra et al., 2025). The system can operate using either observed or forecast weather data and recommends fungicide application within 14 days before or after an infection alert, while maintaining a minimum interval of 14 days between consecutive applications and accounting for pre-harvest intervals as harvest approaches. PedMan was evaluated under commercial grove conditions in the highly susceptible cv. Souri and demonstrated reliable performance in predicting disease risk and supporting effective disease management (Gilat et al., 2026).

The integration of quantitative cultivar resistance into disease forecasting systems provides a promising approach for adapting fungicide recommendations to the inherent susceptibility of the host. Quantitative resistance can be incorporated into disease forecasting and DSS frameworks in several ways, including adjustment of epidemiological or fungicide-risk thresholds, modification of the predicted rate of disease development, or implementation of cultivar-specific rules for initiating and repeating fungicide applications. These concepts have been demonstrated experimentally in several pathosystems. For example, Abuley and Nielsen (2019) incorporated different levels of cultivar resistance to potato early blight (*Alternaria solani*) into the TOMCAST forecasting model. Their results demonstrated that higher TOMCAST thresholds could be successfully applied to more resistant cultivars, whereas lower thresholds were required for susceptible cultivars. A similar principle has been incorporated into late blight DSSs, in which cultivar resistance is considered together with observed and forecast weather conditions, disease development, and fungicide persistence to determine cultivar-specific spray schedules (Small et al., 2015). In the BlightPro DSS, for example, cultivar resistance was explicitly incorporated into disease simulations and field-based evaluations of fungicide scheduling, resulting in fewer applications for moderately resistant cultivars while maintaining effective disease suppression (Small et al., 2015; Wu et al., 2017).

A similar approach is particularly relevant to the management of olive peacock eye disease. Building on these concepts, the PedMan DSS, originally developed and validated for highly susceptible olive cultivars (Ezra et al., 2025; Gilat et al., 2026), can be extended by incorporating cultivar-specific quantitative resistance as an additional component of its decision algorithm. Consequently, highly susceptible cultivars would retain relatively conservative thresholds, whereas moderately susceptible and moderately resistant cultivars would require a greater accumulation of infection risk before a fungicide application is recommended. Such an approach would enable PedMan to integrate pathogen-favorable weather conditions with host susceptibility and thereby provide cultivar-specific spray recommendations, potentially reducing unnecessary fungicide applications while maintaining adequate protection against peacock spot.

The long-term goal of our research was to develop effective and environmentally friendly means for managing peacock eye disease of olive. In the present study, we expanded the PedMan system to enable its reliable application across olive cultivars differing in their susceptibility to *V. oleaginea*. We then evaluated the performance of the expanded system under natural infection conditions in commercial olive groves representing different levels of cultivar susceptibility. Finally, the results from the present study and the parallel report (Gilat et al., 2026) were analyzed together to provide a comprehensive validation of the PedMan system across olive cultivars differing in susceptibility to *V. oleaginea*.

## 2. Materials and Methods

### 2.1. Expanding the criteria used by PedMan to cultivars exhibiting different levels of response to V. oleaginea

The criteria currently used by PedMan to issue an infection alert were developed and evaluated for olive cultivars that are highly susceptible to *V. oleaginea* (Gilat et al., 2026). The objective of the present study was to adapt these criteria for cultivars that are moderately susceptible or moderately resistant to *V. oleaginea* by reducing the probability of issuing infection alerts by approximately 25% and 50%, respectively. To achieve this objective, we used weather data recorded at 11 meteorological stations located in the main olive-growing region of Israel during the 2020/1–2023/4 growing seasons (Gilat et al., 2026). These data were used in simulation runs in which PedMan identified the occurrence and timing of infection alerts during the autumn-early winter (September–December) and spring (March–May) infection periods. In the simulations, the rainfall criterion currently used for highly susceptible cultivars (a rainfall event of ≥15 mm within a single day or accumulated over several consecutive days) was increased to either ≥25 or ≥45 mm. Similarly, the temperature criterion (mean daily minimum temperature during rainy days between 12 and 18 °C) was narrowed to either 13–17 °C or to 14–16 °C. In a previous study (Ygzao et al., 2026a), we found that 83% of *V. oleaginea* conidia germinated within the temperature range of 12–18 °C. Restricting the temperature range to 13–17 °C reduced conidial germination to 65.8% (a 20.7% reduction), whereas restricting it to 14–16 °C reduced germination further, to 40.2% (a 51.5% reduction).

The number of infection alerts generated for each combination of rainfall threshold and temperature range was compared with the number of alerts generated using the criteria for the highly susceptible cultivars, at each weather station. This analysis enabled us to identify the combinations of rainfall and temperature thresholds expected to reduce the frequency of infection alerts by approximately 25% and 50%, corresponding to cultivars with moderate susceptibility and moderate resistance, respectively.

Characterization of the response of the main olive cultivars grown in Israel to *V. oleaginea* was based on the multi-year experience of leading extension officers (Reuven Birger, Mugira Yunis, and Nizar Abed-Al-Ahadi), results from inoculation experiments using detached leaves, and evaluations conducted during the 2022/3 to 2024/5 growing seasons in the Israeli olive germplasm collection at the Volcani Institute in Rishon LeZion, Israel (Gilat et al., unpublished data). Based on these data, cvs. Souri and Nabali were classified as highly susceptible; cvs. Manzanillo, Coratina, Picual and Arbequina as moderately susceptible; cvs. Picholine, Koroneiki and Barnea as moderately resistant; and cvs. Askal, Mission, Kadesh, Ma’alot and Miilya as highly resistant. As for the highly susceptible cultivars (Ezra et al., 2025), the criteria considered when issuing infection alerts for moderately susceptible cultivars were limited to rainfall quantity and minimum temperature range, as specified below. For moderately resistant cultivars, in addition to these criteria, the time of year (autumn, winter, or spring) and the presence of symptomatic leaves during the previous or current growing season were also considered. According to PedMan, spraying is not recommended for cultivars that are highly resistant to *V. oleaginea*.

### 2.2. Evaluating the performance of the revised PedMan system under natural conditions in commercial olive groves

#### 2.2.1. Experimental locations

PedMan recommendations were evaluated in two grove experiments during the 2024/5 growing season and in 22 commercial grove observations during the 2025/6 growing season. The two experiments were conducted in Kfar Ben Nun (31.859262° N, 34.951702° E), located in the Inner Plains region of Israel. Experiment 1 was established in olive trees of cv. Picual, whereas Experiment 2 was conducted in cv. Manzanillo. The commercial observations were carried out at two locations. The first consisted of a 22.5-ha olive grove near Kibbutz Revadim (31.788228° N, 34.827163° E) in the Inner Plains region. The second comprised three olive plots near Kibbutz Yodfat in the Lower Galilee Mountains: Plot 1 (4.5 ha; 32.781290° N, 35.258152° E), Plot 2 (7.5 ha; 32.793788° N, 35.250454° E), and Plot 3 (31.5 ha; 32.788709° N, 35.246459° E). The Revadim grove consisted of 12 subplots (R1–R12), of which six were planted with cv. Coratina, two with cv. Picual, two with cv. Barnea, and two with cv. Askal. Plot 1 at Yodfat consisted of two subplots planted with cv. Coratina (Y1a and Y1b). Plot 2 consisted of three Coratina subplots (Y2a–Y2c). Plot 3 comprised six subplots: two planted with cv. Picual (Y16 and Y17), two with cv. Picholine (Y4 and Y5), and two with cv. Barnea (Y10 and Y11).

At both locations, the olive trees were more than 30 years old and were planted at spacings of 6–7 m within rows and 7 m between rows. Before the initiation of the study, none of the groves had received fungicide treatments against *V. oleaginea*. During the experiments and observations, irrigation, fertilization, pruning, weed and insect control, and harvesting were performed by the growers according to the recommendations of the Israeli Extension Service. Meteorological data required by PedMan, including daily rainfall and minimum air temperature, were obtained from the Israel Meteorological Service and the Ministry of Agriculture and Food Security (www.agrometeorology.co.il). Data were collected from the Nakhshon, Revadim, and Hinaton meteorological stations, which are located within 2 km of the study sites at Kfar Ben Nun, Revadim, and Yodfat, respectively.

#### 2.2.2. Experimental procedures and treatments

The original objective of the 2024/5 field experiments was to evaluate the positive recommendations of PedMan by applying fungicides according to the system’s recommendations. However, during the autumn of the 2024/5 growing season, the minimum temperature during all rainfall events at Kfar Ben Nun remained below 12°C. Consequently, none of the rainfall events met the PedMan criteria for an infection event, and the system did not recommend fungicide application. To evaluate the reliability of PedMan’s negative recommendation (i.e., withholding fungicide applications), fungicides were applied following a rainfall event exceeding 15 mm that occurred on 26 December 2024 (Figure 1A), despite the absence of a PedMan recommendation. Accordingly, the experiments included two treatments: (i) **Control**, in which trees were sprayed with fungicides against *V. oleaginea* despite PedMan recommending no treatment, and (ii) **PedMan**, in which trees were left unsprayed in accordance with the system’s recommendation.

**Figure 1.**
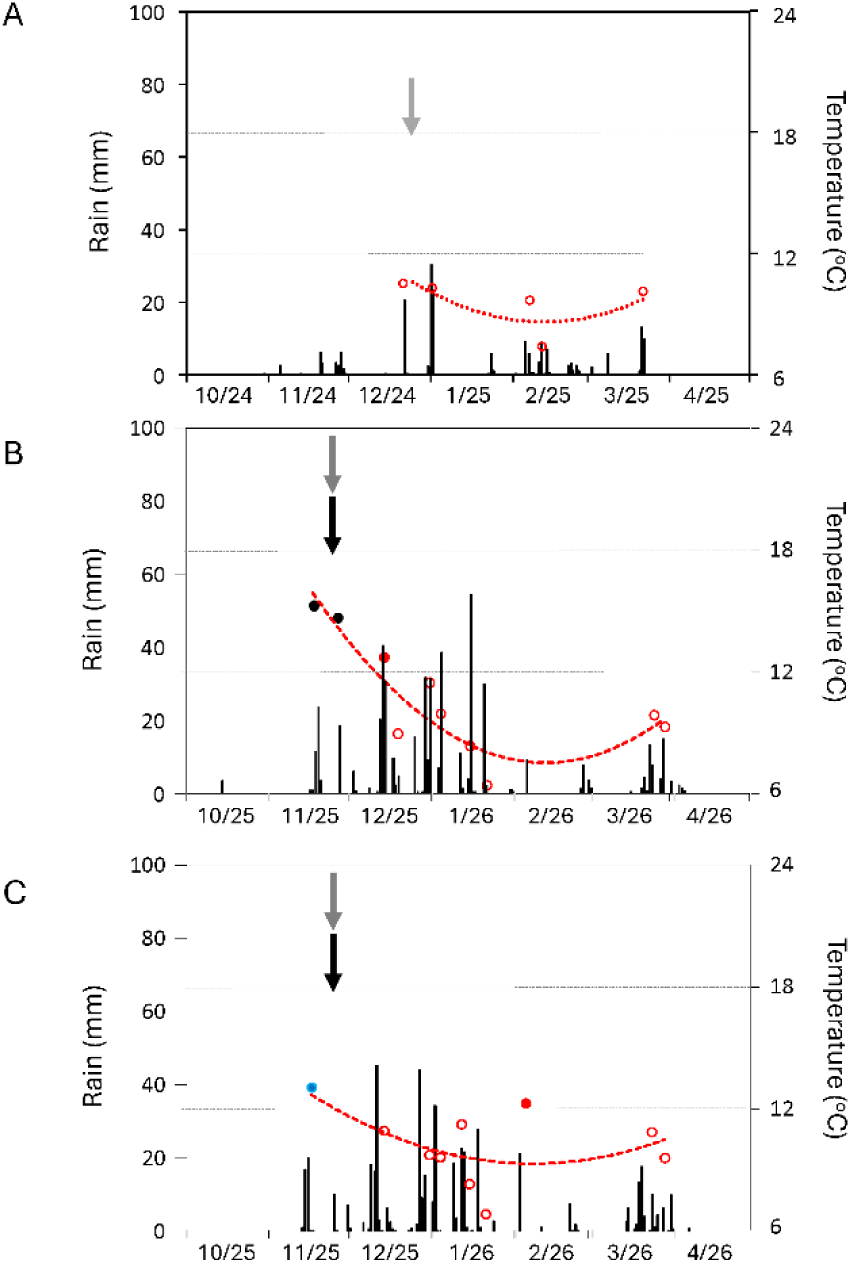
Weather conditions recorded at meteorological stations located near the experimental sites during the 2024/5 growing season at Kfar Ben Nun (**A**) and near the observation sites at Revadim (**B**) and Yodfat (**C**) during the 2025/6 growing season. Bars represent daily rainfall. Circles indicate the average minimum temperature during rainfall events exceeding 15 mm. Black filled circles denote rainfall events that met the PedMan criteria for infection in highly susceptible, moderately susceptible, and moderately resistant cultivars. Blue filled circles denote rainfall events that met the PedMan criteria for infection in highly susceptible and moderately susceptible cultivars, whereas red filled circles denote rainfall events that met the PedMan criteria for infection only in highly susceptible cultivars. Open circles indicate rainfall events that did not meet the PedMan criteria for infection for all cultivar response classes. Red dashed lines show the seasonal trend in minimum temperature. Black arrows indicate fungicide applications made according to PedMan recommendations, whereas gray arrows indicate applications made contrary to PedMan recommendations.

The experiments were arranged in a randomized complete block design with two treatments, four replicates per treatment, and one tree per replicate. The fungicides used were Skipper (difenoconazole, 250 g L⁻¹ EC; Tapazol Chemical Works Ltd., Beit Shemesh, Israel) and Nechostan (190 g L⁻¹ copper as 340 g L⁻¹ tribasic copper sulfate; Nufarm, distributed in Israel by Adama-Agan, Ashdod, Israel). Both fungicides are registered in Israel for the control of peacock eye and are widely used in commercial olive production. They were applied as a tank mixture at concentrations of 0.06% (Skipper) and 0.3% (Nechostan). Fungicides were applied with a handgun sprayer using 15 L of spray solution per tree.

#### 2.2.3. Observation procedures and treatments

As in the field experiments, each of the 22 commercial grove observations consisted of two treatments: **Control** and **PedMan**. The assignment of treatments depended on the recommendation generated by PedMan (Table 1). When PedMan recommended fungicide application (positive recommendation), trees assigned to the PedMan treatment were sprayed, whereas those assigned to the control treatment remained unsprayed to evaluate the benefit of the recommended application. Conversely, when PedMan recommended that no fungicide should be applied (negative recommendation), trees assigned to the PedMan treatment were left unsprayed, whereas those assigned to the control treatment were sprayed to evaluate the validity of withholding the application.

**Table 1.** Details about the observations carried out in the 2025/6 growing season in commercial olive groves to evaluating the performance of PedMan recommendations in cultivars with dissimilar responses to *V. oleaginea*.

| Location | Cultivar |  | Subplot | Previous | PedMan | Treatment<sup>z</sup> |  |
|---|---|---|---|---|---|---|---|
|  | Name | response<sup>y</sup> | code | infection | Recomm. | Control | PedMan |
| Revadim | Coratina | MS | R1 | Yes | To spray | Not sprayed | Sprayed |
|  |  |  | R3 | Yes | To spray | Not sprayed | Sprayed |
|  |  |  | R6 | Yes | To spray | Not sprayed | Sprayed |
|  |  |  | R8 | Yes | To spray | Not sprayed | Sprayed |
|  |  |  | R10 | Yes | To spray | Not sprayed | Sprayed |
|  |  |  | R11 | Yes | To spray | Not sprayed | Sprayed |
|  | Picual | MS | R4 | Yes | To spray | Not sprayed | Sprayed |
|  |  |  | R7 | Yes | To spray | Not sprayed | Sprayed |
|  | Barnea | MR | R2 | No | Not to spray | Sprayed | Not sprayed |
|  |  |  | R5 | No | Not to spray | Sprayed | Not sprayed |
|  | Askal | HR | R9 | Yes | Not to spray | Sprayed | Not sprayed |
|  |  |  | R12 | Yes | Not to spray | Sprayed | Not sprayed |
| Yodfat | Coratina | MS | Y1a | No | To spray | Not sprayed | Sprayed |
|  |  |  | Y1b | No | To spray | Not sprayed | Sprayed |
|  |  |  | Y2a | No | To spray | Not sprayed | Sprayed |
|  |  |  | Y2b | No | To spray | Not sprayed | Sprayed |
|  |  |  | Y2c | No | To spray | Not sprayed | Sprayed |
|  | Picual | MS | Y16-17 | Yes | To spray | Not sprayed | Sprayed |
|  | Picholine | MR | Y4 | No | Not to spray | Sprayed | Not sprayed |
|  |  |  | Y5 | No | Not to spray | Sprayed | Not sprayed |
|  | Barnea | MR | Y10 | No | Not to spray | Sprayed | Not sprayed |
|  |  |  | Y11 | No | Not to spray | Sprayed | Not sprayed |

<sup>y</sup>Cultivar resistance class: MS = moderately susceptible; MR = moderately resistant; HR = highly resistant.
<sup>z</sup>in cases where PedMan recommendation was 'not to spray' the sprays were applied in the control treatment to evaluate the negative accuracy of PedMan (applying sprays in contrast to system's recommendation).

At the Revadim site, the grower sprayed one row (30–40 trees) in each of the 12 subplots on 19 November 2025 (Figure 1B). Depending on the PedMan recommendation for each subplot, the sprayed row served either as the PedMan treatment (when spraying was recommended) or as the control treatment (when no spray was recommended). The remaining unsprayed trees constituted the corresponding comparison treatment. At the Yodfat site, PedMan recommended fungicide application in the Coratina and Picual subplots. Accordingly, three rows of trees in each subplot were left unsprayed as the control treatment, whereas the remaining trees were sprayed according to PedMan recommendations. In contrast, PedMan did not recommend fungicide application in the Picholine and Barnea subplots. Therefore, three rows of trees in each subplot were sprayed contrary to the PedMan recommendation (control treatment), whereas the remaining trees were left unsprayed (PedMan treatment). Fungicide applications at both sites were made on 19 November 2025 (Figure 1B and C). In all observations, growers applied a tank mixture of Skipper (0.06%) and Nechostan (0.3%) using a Degania axial fan sprayer at a spray volume of 2,000 L ha⁻¹.

#### 2.2.4. Evaluation of PedMan recommendations in the observations

The impact of peacock eye disease on tree health was assessed by estimating leaf abscission. Leaf abscission was selected as the response variable because it integrates the cumulative effect of repeated infections on tree health and is the principal cause of yield reduction associated with peacock eye disease (Graniti, 1993; Buonaurio et al., 2023). Leaf abscission (expressed as %) was estimated as previously described (Ygzao et al. 2026c; Gilat et al., 2026). Briefly, each tree canopy was visually assessed from four directions, and the percentage of leaf abscission was estimated for each orientation. A score of 0% indicated a full, healthy canopy with no visible leaf loss; A score of 50% indicated that approximately half of the leaves had abscised and the canopy appeared noticeably transparent, and A score of 100% indicated complete defoliation. The four directional estimates were averaged to obtain a single value for each tree. In the field experiments, leaf abscission was assessed for each experimental tree (one tree per replicate) every 2-3 weeks starting at early January and ending at late May, 2026. In the commercial grove observations, 20 trees were evaluated for each treatment within each subplot. Assessments were conducted in April or May, corresponding to the peak of the second annual peacock eye disease episode (Ygzao et al., 2026c).

Leaf abscission in olive trees may also result from other biotic and abiotic factors, including other foliar pathogens, insect pests, drought, waterlogging, salinity, nutrient imbalance, and temperature extremes (Martin and Sibbett, 2005). Nevertheless, the differences in leaf abscission observed between fungicide-treated and untreated trees in the present study were attributed primarily to peacock eye disease, as all other cultural practices were identical between treatments within each experiment or observation. For each experiment and observation, control efficacy (CE, %) was calculated by comparing the incidence of leaf abscission in the PedMan treatment (*LA_t_*) with the incidence of leaf abscission in the control treatment (*LA_c_*) according to the following formula:

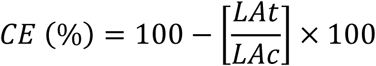

### 2.3. Data analyses

In accordance with the two types of operational decisions generated by PedMan, the data were analyzed separately for cases in which fungicide application was recommended (spray recommendations; cvs. Coratina and Picual) and cases in which fungicide application was not recommended (no-spray recommendations; cvs. Manzanillo, Barnea, Picholine, and Askel). The field observations were analyzed using two complementary approaches. First, differences in the incidence of leaf abscission between treated and untreated trees were assessed using a two-sided Student’s *t*-test assuming equal variances. Second, to account for potential spatial pseudo replication among trees within rows and the unbalanced structure of the observations, a linear mixed-effects model (LMM) was fitted using restricted maximum likelihood (REML) estimation. The LMM incorporated treatment as a fixed effect and row as a random effect, thereby accounting for the non-independence of observations within rows. Subplot was incorporated as a random intercept effect to serve as a spatial block factor. Fixed effects in both models comprised ‘location’, ‘cultivar’ and ‘treatment’, and the ‘treatment’ × ‘cultivar’ interaction term. Statistical analyses were executed using JMP software (Version 16; SAS Institute, Cary, NC). The linear mixed-effects model for each independent validation phase was expressed using the following algebraic equation:

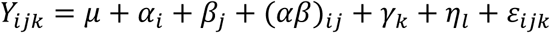

Where:

- *Y_ijk_* represents the observed incidence of leaf abscission (%) quantified at the row level.
- *μ* is the overall experimental mean baseline intercept.
- *α_i_*is the fixed effect of the *i* -th Treatment (where *i* = PedMan recommendation vs. Control).
- *β_j_* is the fixed effect of the *j* -th Cultivar.

`o` For the positive accuracy phase: *j* = Coratina, Picual.
`o` For the negative accuracy phase: *j* = Barnea, Fishulin, Askal.
- (*αβ*)*_ij_* is the fixed interaction effect between the *i* -th Treatment and the *j* -th Cultivar, evaluating the variance in treatment efficacy across different cultivars.
- *γ_k_* is the fixed environmental covariate effect of the *k* -th location *k* = (Revadim, Yodfat) to account for regional macroclimatic baseline variation.
- *η_l_* is the random intercept effect of the *l* -th treatment, acting as a spatial blocking factor to pair the treatment and control rows within each subplot. Normality assumptions were tested and verified to be independently distributed: 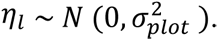
- *ε_ijk_* is the leftover residual random error term representing within-plot row variance, Normality assumption was tested and verified to be independently distributed: 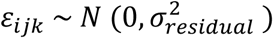

### 2.4. Evaluation of PedMan recommendations across olive cultivars differing in susceptibility to V. oleaginea

PedMan recommendations were implemented in 33 field experiments and observations in total. Of which, 9 were presented in the parallel report (Gilat at al., 2026) and 24 in the current report. The performance of the system was evaluated using a binary classification framework based on the statistical evidence obtained from each field trial or observation. The 33 evaluations were classified according to whether the experimental results supported the corresponding PedMan recommendation. When PedMan recommended fungicide application, the recommendation was considered as correct when a statistically significant difference in leaf abscission was detected between trees receiving the fungicide treatment recommended by PedMan and untreated control trees. Conversely, when PedMan recommended no fungicide application, the recommendation was considered as correct when no statistically significant difference in leaf abscission was detected between trees left untreated according to the PedMan recommendation and trees that received fungicide despite the no-spray recommendation. Thus, a statistically significant treatment effect supported a spray recommendation, whereas the absence of a statistically significant treatment effect supported a no-spray recommendation.

The resulting decisions were summarized in a contingency table, where the recommendation to spray defined as the positive class and withholding spray as the negative class. Outcomes were partitioned into true positive (TP), true negative (TN), false positive (FP), and false negative (FN). Because classification accuracy alone is insufficient for evaluating performance on potentially imbalanced binary tasks (Altman et al., 1994; Foody, 2023; Chicco and Jurman, 2023), we jointly evaluated overall accuracy, sensitivity, specificity, positive predictive value (PPV) and negative predictive value (NPV). Overall accuracy represents the proportion of all predictions that were correct, including both correct positive and correct negative predictions; sensitivity represents the proportion of evaluations in which the statistical analysis supported fungicide application and PedMan correctly recommended spraying; specificity represents the proportion of evaluations in which the statistical analysis did not support fungicide application and PedMan correctly recommended no spraying. PPV and NPV were considered jointly because they characterize different aspects of binary classification performance and accuracy alone does not fully describe the performance of a classification system (Altman et al., 1994; Foody, 2023; Chicco and Jurman, 2023). These performance metrics were calculated as follows:

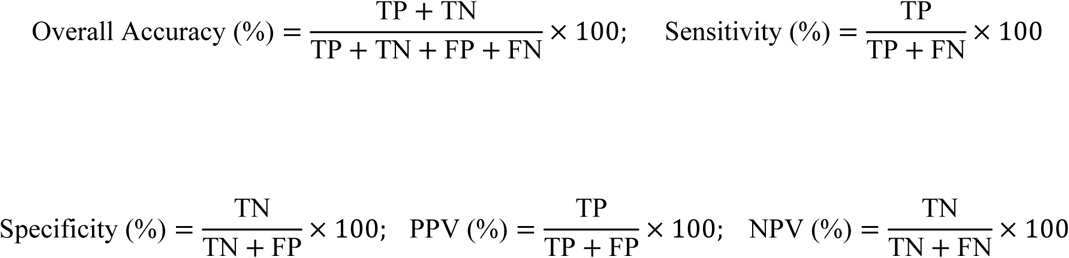

As complementary measures of overall classification performance, balanced accuracy, the F1 score, and the Matthews correlation coefficient (MCC) were calculated from the contingency table. Balanced accuracy quantifies the average ability of PedMan to correctly classify both disease-risk and non-disease-risk cases, giving equal weight to the sensitivity and specificity of the predictions. F1 score quantifies the balance between PedMan’s precision and sensitivity, reflecting its ability to correctly identify disease-risk cases while minimizing both missed cases and false-positive predictions. MCC quantifies the overall agreement between PedMan predictions and the observed disease-management outcomes, accounting for both correct and incorrect positive and negative classifications. It was included because it incorporates all four cells of the contingency table and provides a single summary measure of agreement between the predicted and reference classifications (Chicco and Jurman, 2023). Because of the relatively small validation dataset and the occurrence of observed proportions equal to 100%, two-sided 95% confidence intervals for sensitivity, specificity, PPV, and NPV were calculated using exact binomial confidence limits. The 95% confidence interval for overall accuracy was calculated using the Wilson score method. All confidence intervals were two-sided and calculated at the 95% confidence level.

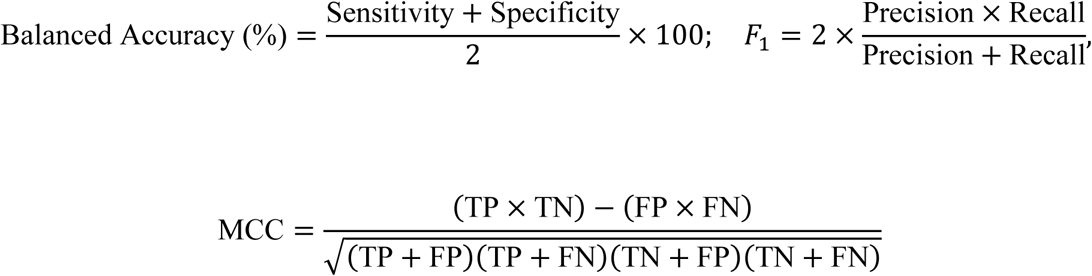

## 3. Results

### 3.1. Expanding the criteria used by PedMan to cultivars exhibiting different levels of response to V. oleaginea

Data collected over four growing seasons from 11 meteorological stations representing all olive-growing regions in Israel were used to perform simulation runs aimed at defining rainfall and temperature criteria for cultivars with different levels of susceptibility to *V. oleaginea*. First, only the temperature criterion was calibrated; Narrowing the thermal window used by the system for the highly susceptible cultivars (12–18 °C) to 13–17 °C reduced the mean number of infection alerts issued by the system by 25.1%, whereas further narrowing it to 14–16 °C, resulted in a 51.3% reduction (Figure 2A and B). Next, only the rainfall threshold was modified. Increasing the rainfall threshold used for highly susceptible cultivars (≥15 mm) to ≥25 or ≥45 mm, reduced the mean number of infection alerts by 34.9% and 55.1%, respectively (Figure 2C and D). Finally, both the temperature and rainfall criteria were modified simultaneously. Combining a rainfall threshold of ≥25 mm with a temperature range of 13–17 °C reduced the mean number of infection alerts by 54.0%, whereas combining a rainfall threshold of ≥45 mm with a temperature range of 14–16°C reduced the number of alerts by 84.1% (Figure 2E and F).

**Figure 2.**
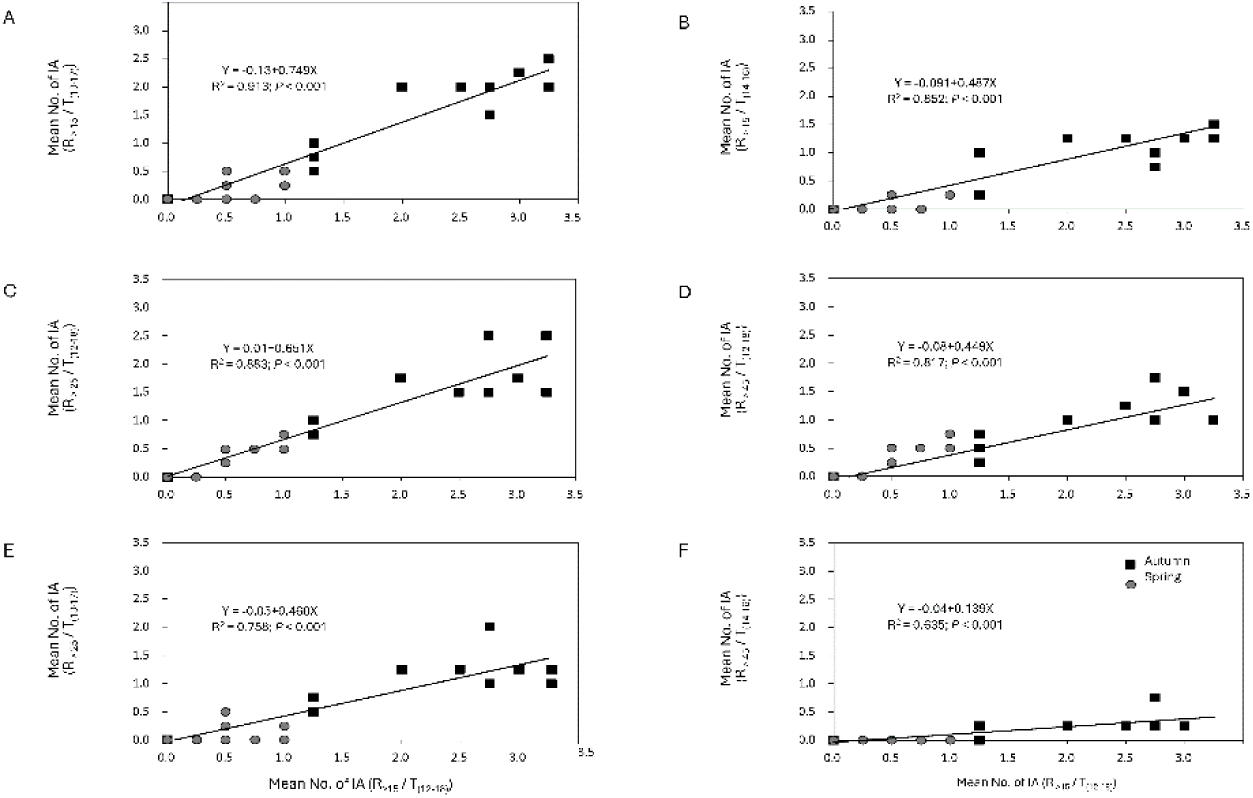
Sensitivity analysis of the thresholds used by PedMan to generate infection alerts (IA). The mean number of infection alerts issued during autumn (black squares) and spring (gray circles) over four growing seasons (2020/1–2023/4) at 11 weather stations was calculated using the standard PedMan thresholds (R_>15_/T_12–18_) and compared with the mean number of alerts generated using different rain and temperature definitions.

Because our objective was to reduce the probability of issuing infection alerts by approximately 25% and 50% for moderately susceptible and moderately resistant cultivars, respectively, only the temperature criterion was modified. Accordingly, a temperature range of 13–17°C was adopted as the threshold for issuing infection alerts for moderately susceptible cultivars, whereas a narrower range of 14–16°C was adopted for moderately resistant cultivars.

### 3.2. Evaluating the performance of the revised PedMan system under natural conditions in commercial olive groves

#### 3.2.1. Evaluating the positive predictive performance of PedMan recommendations

The positive predictive performance of PedMan recommendations was evaluated during the 2025/6 growing season in 14 observations carried out in commercial olive groves planted with cultivars Coratina and Picual. Comparing the differences between the treatments in each observation revealed that following PedMan recommendation to spray reduced leaf abscission significantly (*P* ≤ 0.05) in all cases (Table 2). The entire data set was then analyzed using the Linear Mixed Model. The fixed effects (’location’, ‘cultivar’ and ‘treatment’) explained 80.7% of the total defoliation variance (Marginal *R^2^* = 0.807), while the total model (accounting for the random subblock effect) profiles explained 96.7% of the variance (Conditional *R^2^* = 0.967). The random spatial subplot effect accounted for 72.7% of the remaining unresolved variance components (Table 3), confirming strong environmental spatial dependency among trees within the same grove subplot.

**Table 2.**
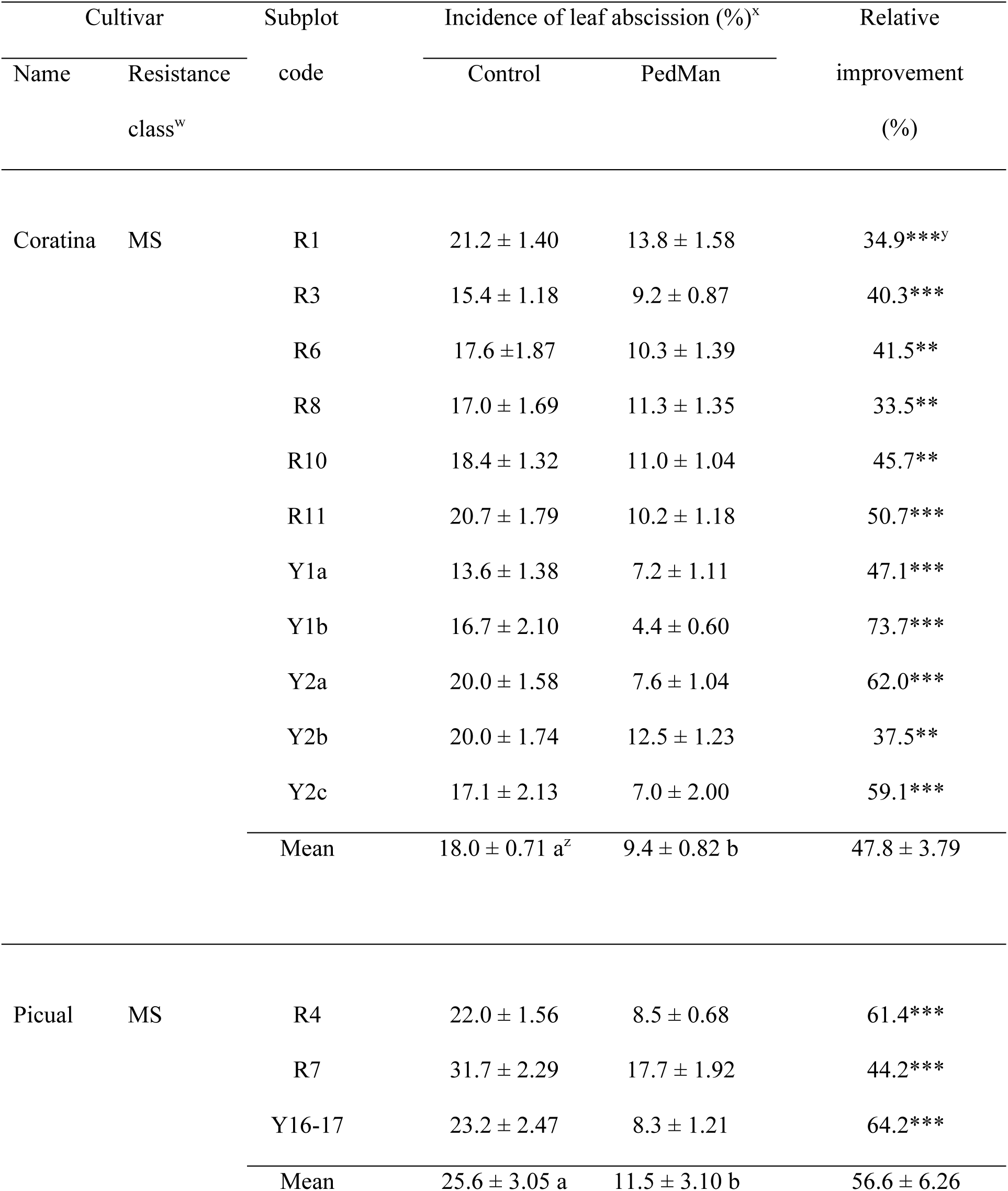
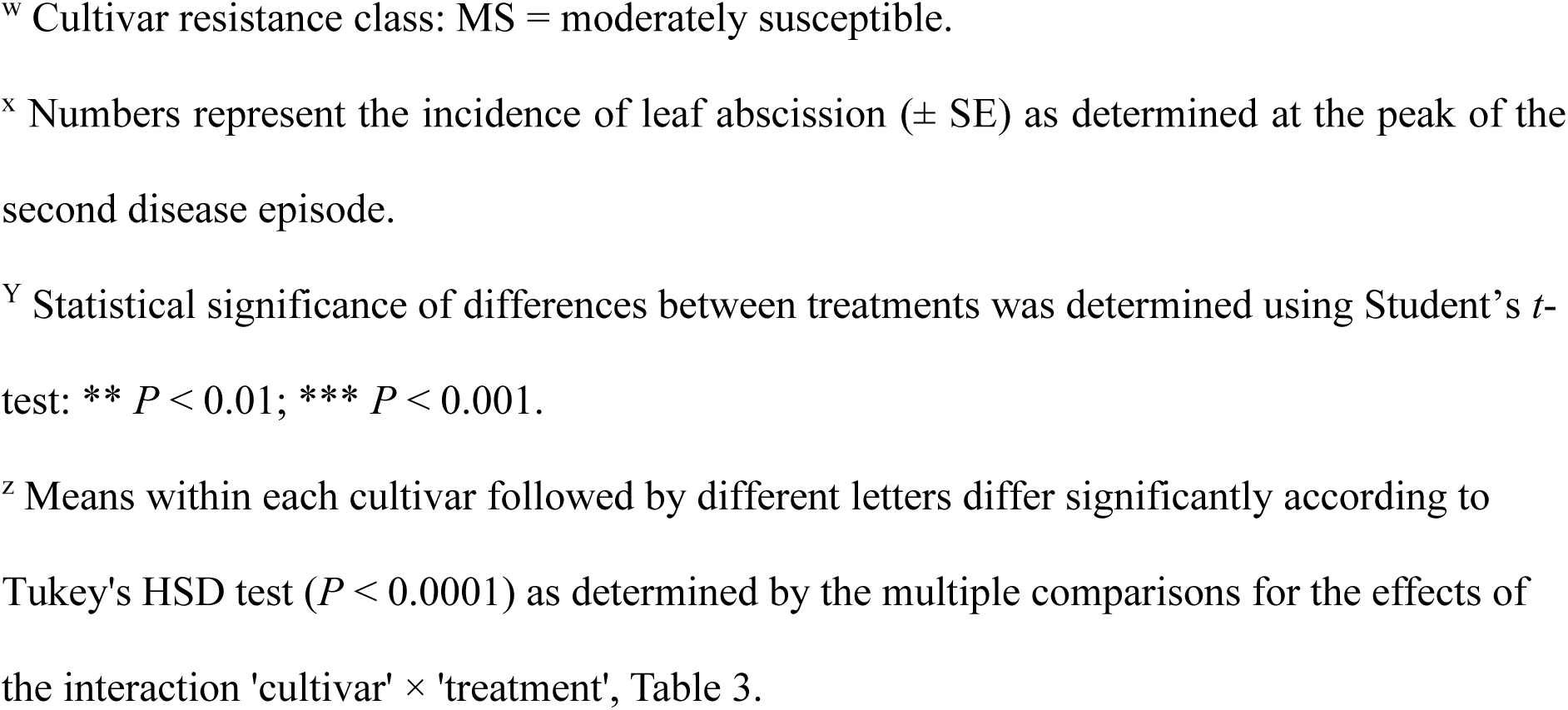
Evaluating the positive predictive performance of PedMan recommendations (applying sprays in accordance to system’s recommendation) in cultivars that were classified as moderately susceptible to *V. oleaginea*.

| Cultivar |  | Subplot code | Incidence of leaf abscission (%)<sup>x</sup> |  | Relative improvement (%) |
|---|---|---|---|---|---|
| Name | Resistance class<sup>w</sup> |  | Control | PedMan |  |
| Coratina | MS | R1 | 21.2 ± 1.40 | 13.8 ± 1.58 | 34.9***<sup>y</sup> |
|  |  | R3 | 15.4 ± 1.18 | 9.2 ± 0.87 | 40.3*** |
|  |  | R6 | 17.6 ±1.87 | 10.3 ± 1.39 | 41.5** |
|  |  | R8 | 17.0 ± 1.69 | 11.3 ± 1.35 | 33.5** |
|  |  | R10 | 18.4 ± 1.32 | 11.0 ± 1.04 | 45.7** |
|  |  | R11 | 20.7 ± 1.79 | 10.2 ± 1.18 | 50.7*** |
|  |  | Y1a | 13.6 ± 1.38 | 7.2 ± 1.11 | 47.1*** |
|  |  | Y1b | 16.7 ± 2.10 | 4.4 ± 0.60 | 73.7*** |
|  |  | Y2a | 20.0 ± 1.58 | 7.6 ± 1.04 | 62.0*** |
|  |  | Y2b | 20.0 ± 1.74 | 12.5 ± 1.23 | 37.5** |
|  |  | Y2c | 17.1 ± 2.13 | 7.0 ± 2.00 | 59.1*** |
|  |  | Mean | 18.0 ± 0.71 a<sup>z</sup> | 9.4 ± 0.82 b | 47.8 ± 3.79 |
| Picual | MS | R4 | 22.0 ± 1.56 | 8.5 ± 0.68 | 61.4*** |
|  |  | R7 | 31.7 ± 2.29 | 17.7 ± 1.92 | 44.2*** |
|  |  | Y16-17 | 23.2 ± 2.47 | 8.3 ± 1.21 | 64.2*** |
|  |  | Mean | 25.6 ± 3.05 a | 11.5 ± 3.10 b | 56.6 ± 6.26 |

<sup>w</sup> Cultivar resistance class: MS = moderately susceptible.
<sup>x</sup> Numbers represent the incidence of leaf abscission (± SE) as determined at the peak of the second disease episode.
<sup>Y</sup> Statistical significance of differences between treatments was determined using Student’s *t*-test: \*\* *P* < 0.01; \*\*\* *P* < 0.001.
<sup>z</sup> Means within each cultivar followed by different letters differ significantly according to Tukey's HSD test (*P* < 0.0001) as determined by the multiple comparisons for the effects of the interaction 'cultivar' × 'treatment', Table 3.

**Table 3.**
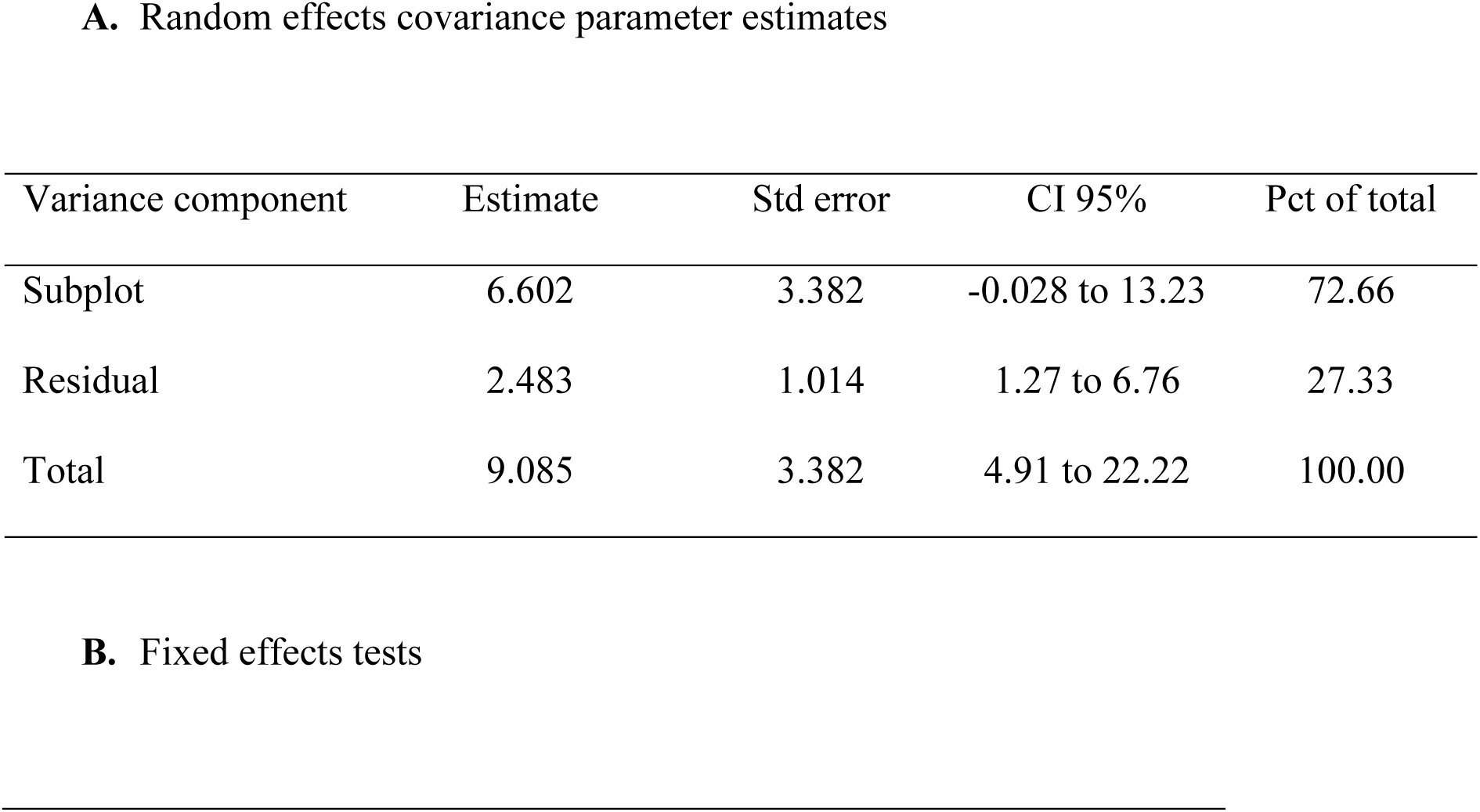
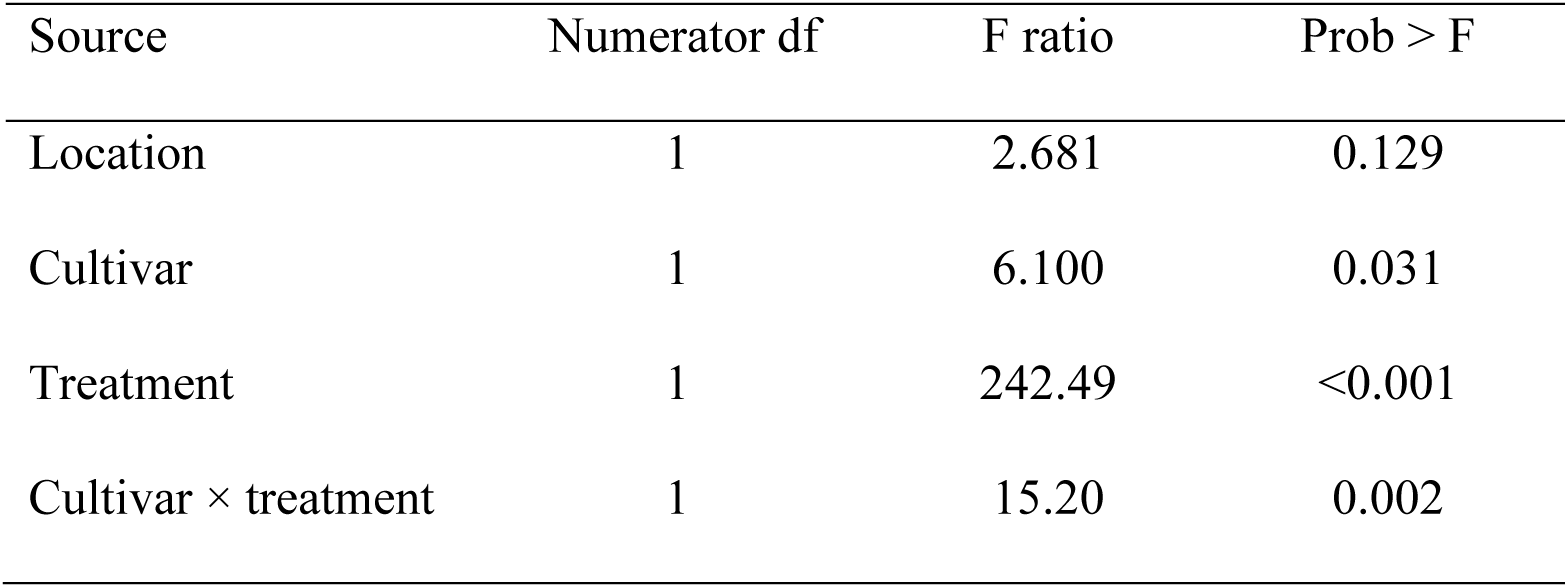
Linear mixed-model analysis of factors affecting leaf abscission in the observation carried out for evaluating the positive predictive performance of PedMan recommendations (applying sprays in accordance to system’s recommendation) in cultivars that were classified as moderately susceptible to *V. oleaginea*.

| **A.** Random effects covariance parameter estimates |  |  |  |  |
|---|---|---|---|---|
| Variance component | Estimate | Std error | CI 95% | Pct of total |
| Subplot | 6.602 | 3.382 | -0.028 to 13.23 | 72.66 |
| Residual | 2.483 | 1.014 | 1.27 to 6.76 | 27.33 |
| Total | 9.085 | 3.382 | 4.91 to 22.22 | 100.00 |
| **B.** Fixed effects tests |  |  |  |  |
| Source | Numerator df | F ratio | Prob > F |  |
| Location | 1 | 2.681 | 0.129 |  |
| Cultivar | 1 | 6.100 | 0.031 |  |
| Treatment | 1 | 242.49 | <0.001 |  |
| Cultivar × treatment | 1 | 15.20 | 0.002 |  |

’Treatment’ exerted strongest influence on leaf abscission (*F* = 242.49; *P* < 0.001), while ‘cultivar’ also significantly affected leaf abscission (*F* = 6.10; *P* = 0.031). In addition, a significant ‘cultivar’ × ‘treatment’ interaction (*F* = 15.20; *P* = 0.002) indicated that the magnitude of the treatment effect differed between cultivars. Because the ‘cultivar’ × ‘treatment’ interaction was significant, treatment effects were subsequently sliced by cultivar *via* post-hoc Least-Squares means contrasts for each cultivar. For cv. Coratina, PedMan-guided spraying significantly suppressed premature leaf abscission (*P* < 0.0001), reducing leaf abscission from 18.0 to 9.4%, a 47.8% reduction. For cv. Picual, PedMan-guided spraying reduced mean leaf abscission from 25.6 to 11.5%, corresponding to a 56.6% improvement (*P* < 0.001). These field observations therefore confirmed the positive predictive performance of PedMan spray recommendations under commercial grove conditions.

#### 3.2.2. Evaluating the negative predictive performance of PedMan recommendations

The negative predictive performance of PedMan recommendations was evaluated in two grove experiments conducted during the 2024/5 growing season and in eight field observations carried out during the 2025/6 growing season in commercial olive groves planted with the cultivars Picual, Manzanillo, Barnea, Picholine and Askal. In the grove experiment conducted with cv. Picual, withholding fungicide applications according to the PedMan recommendation had no adverse effect on leaf abscission throughout the growing season. Differences in leaf abscission between the two treatments remained insignificant throughout the season. Furthermore, the incidence of leaf abscission in the last assessment date in both treatments did not differ significantly (*P* ≥ 0.05) from the incidence in the first assessment (Figure 3A). These results indicate that fungicide application was unnecessary and confirming the negative predictive performance of the PedMan recommendation. In contrast, in the grove experiment conducted with cv. Manzanillo, trees sprayed despite the PedMan recommendation not to spray exhibited significantly lower leaf abscission during April and May, corresponding to the peak of the second disease episode. Similarly, the incidence of leaf abscission in the last two assessment dates in the PedMan treatment differ significantly (*P* ≤ 0.05) from the incidence in the first assessment. Differences in the incidence of leaf abscission between these date for the control treatment were insignificant (Figure 3B). These results indicate that, under these experimental conditions, the PedMan recommendation not to spray was incorrect.

**Figure 3.**
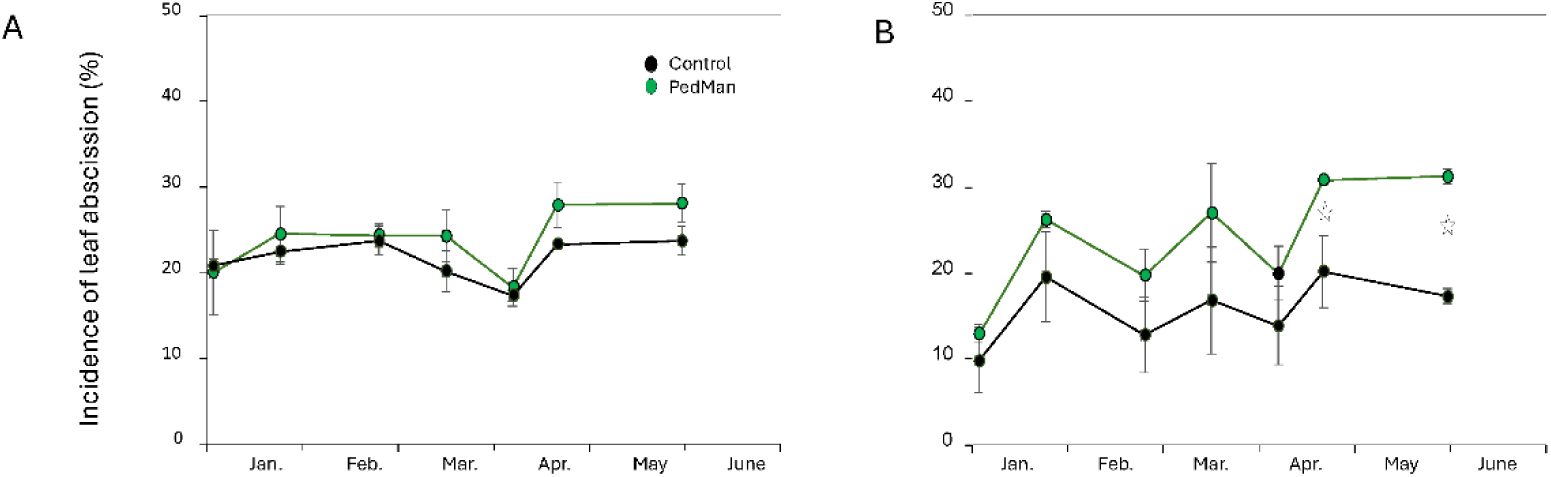
Seasonal progress of leaf abscission in two grove experiments conducted during the 2024/25 growing season to evaluate the negative predictive performance of PedMan recommendations. In the control treatment, fungicides were applied despite the PedMan recommendation not to spray, whereas no fungicides were applied in the PedMan treatment. (**A**) cv. Picual; (**B**) cv. Manzanillo. Vertical bars represent the standard error of the mean. Asterisks indicate significant differences between treatments at the corresponding sampling date (*P* < 0.05).

The negative predictive performance of PedMan recommendations was further evaluated under commercial grove conditions using eight independent field observations. Comparing the differences between the treatments in each observation revealed that following PedMan recommendation not to spray did not affect leaf abscission significantly (*P* ≤ 0.05) for cultivars Barnea and Picholine, but the differences between the treatments were significant in the two observations of cv. Askal (Table 4). When the entire data set was analyzed using the Linear Mixed Model, comparing the differences between the treatments in each observation revealed that following PedMan recommendation to spray reduced leaf abscission significantly (*P* ≤ 0.01) in all cases. The fixed effects of the Linear Mixed Model (’location’, ‘cultivar’ and ‘treatment’) explained 87.6% while the total model (accounting for the random subplot) profiles captured 98.3% of the variance. The random spatial subplot effect accounted for 79.5% of the remaining unresolved variance components, solidifying the equivalent result in the positive recommendations (Table 5).

**Table 4.**
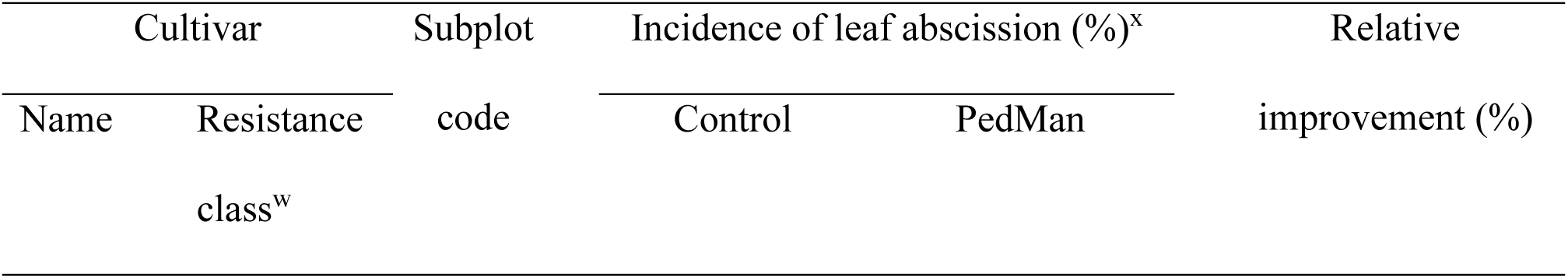
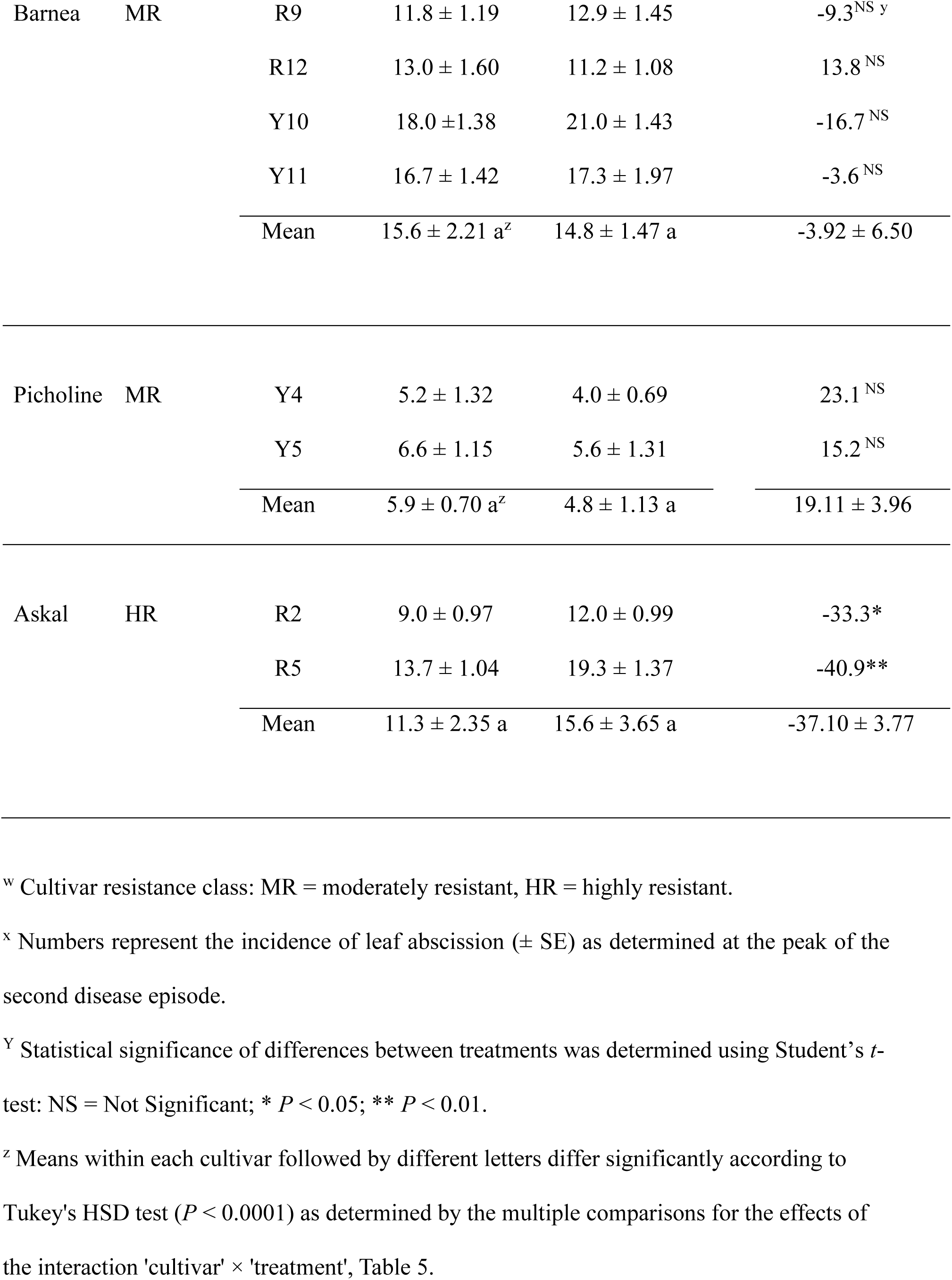
Evaluating the negative predictive performance of PedMan (applying sprays in contrast to system’s recommendation) in cultivars that were classified as moderately of highly resistant to *V. oleaginea*.

| Cultivar |  | Subplot code | Incidence of leaf abscission (%)<sup>x</sup> |  | Relative improvement (%) |
|---|---|---|---|---|---|
| Name | Resistance class<sup>w</sup> |  | Control | PedMan |  |
| Barnea | MR | R9 | 11.8 ± 1.19 | 12.9 ± 1.45 | -9.3<sup>NS y</sup> |
|  |  | R12 | 13.0 ± 1.60 | 11.2 ± 1.08 | 13.8<sup>NS</sup> |
|  |  | Y10 | 18.0 ±1.38 | 21.0 ± 1.43 | -16.7<sup>NS</sup> |
|  |  | Y11 | 16.7 ± 1.42 | 17.3 ± 1.97 | -3.6<sup>NS</sup> |
|  |  | Mean | 15.6 ± 2.21 a<sup>z</sup> | 14.8 ± 1.47 a | -3.92 ± 6.50 |
| Picholine | MR | Y4 | 5.2 ± 1.32 | 4.0 ± 0.69 | 23.1<sup>NS</sup> |
|  |  | Y5 | 6.6 ± 1.15 | 5.6 ± 1.31 | 15.2<sup>NS</sup> |
|  |  | Mean | 5.9 ± 0.70 a<sup>z</sup> | 4.8 ± 1.13 a | 19.11 ± 3.96 |
| Askal | HR | R2 | 9.0 ± 0.97 | 12.0 ± 0.99 | -33.3* |
|  |  | R5 | 13.7 ± 1.04 | 19.3 ± 1.37 | -40.9** |
|  |  | Mean | 11.3 ± 2.35 a | 15.6 ± 3.65 a | -37.10 ± 3.77 |

<sup>w</sup> Cultivar resistance class: MR = moderately resistant, HR = highly resistant.
<sup>x</sup> Numbers represent the incidence of leaf abscission (± SE) as determined at the peak of the second disease episode.
<sup>Y</sup> Statistical significance of differences between treatments was determined using Student’s *t*-test: NS = Not Significant; \* *P* < 0.05; \*\* *P* < 0.01.
<sup>z</sup> Means within each cultivar followed by different letters differ significantly according to Tukey's HSD test (*P* < 0.0001) as determined by the multiple comparisons for the effects of the interaction 'cultivar' × 'treatment', Table 5.

**Table 5.**
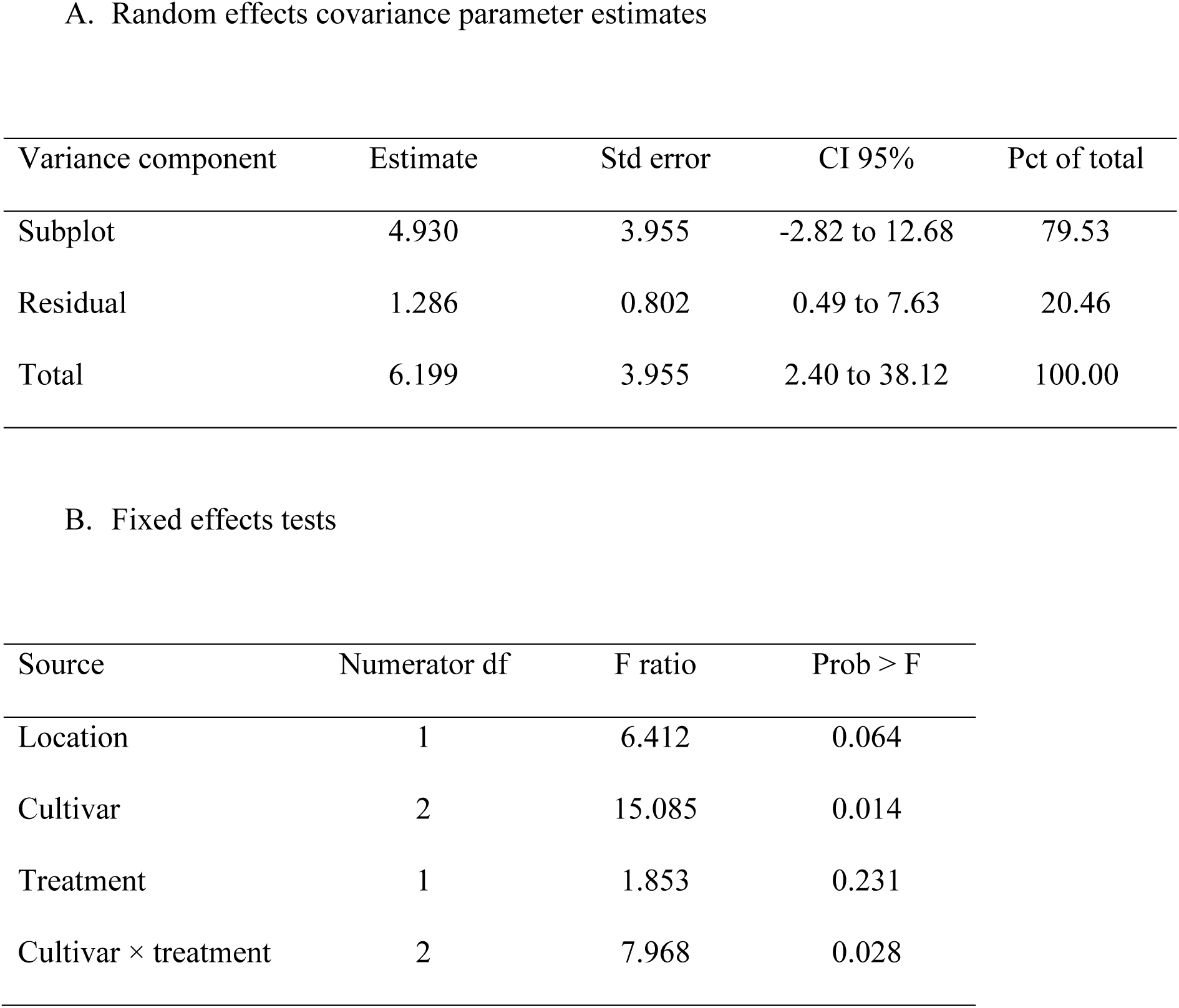
Random and fixed effects of the mixed model used for estimating the parameters affecting leaf abscission in the observation carried out for evaluating the negative predictive performance of PedMan recommendations (applying sprays in contrast to system’s recommendation) in cultivars that were classified as moderately susceptible to *V. oleaginea*.

| A. Random effects covariance parameter estimates |  |  |  |  |
| Variance component | Estimate | Std error | CI 95% | Pct of total |
|---|---|---|---|---|
| Subplot | 4.930 | 3.955 | -2.82 to 12.68 | 79.53 |
| Residual | 1.286 | 0.802 | 0.49 to 7.63 | 20.46 |
| Total | 6.199 | 3.955 | 2.40 to 38.12 | 100.00 |
| B. Fixed effects tests |  |  |  |  |
| Source | Numerator df | F ratio | Prob > F |  |
| Location | 1 | 6.412 | 0.064 |  |
| Cultivar | 2 | 15.085 | 0.014 |  |
| Treatment | 1 | 1.853 | 0.231 |  |
| Cultivar × treatment | 2 | 7.968 | 0.028 |  |

’Cultivar’ significantly affected premature leaf abscission (*F* = 15.09; *P* = 0.013). In contrary, neither ‘location’ (*P* = 0.0645) nor ‘treatment’ (*P* = 0.2315) had a significant overall effect. However, the significant ‘cultivar’ × ‘treatment’ interaction (*F* = 7.97; *P* = 0.027) indicated that host responsiveness to withheld versus applied fungicide sprays varied across different cultivars. Therefore, treatment effects were evaluated separately for each cultivar. For all three cultivars evaluated (Barnea, Picholine, and Askal), applying fungicides despite the PedMan recommendation not to spray did not significantly reduce leaf abscission (*P* > 0.05). Mean leaf abscission remained statistically similar between treated and untreated plots in all cultivars (Table 5). These results demonstrate that fungicide application contrary to the PedMan recommendation provided no measurable benefit, thereby confirming the negative predictive performance of the PedMan decision-support system under commercial grove conditions.

### 3.3. Validation of PedMan recommendations across olive cultivars differing in susceptibility to V. oleaginea

Across the 33 field trials and observations, PedMan provided 30 correct recommendations and three incorrect ones (Table 6), corresponding to an overall accuracy of 90.9% (95% CI: 76.8–96.9%). Of the 23 evaluations in which the statistical analysis supported fungicide application, PedMan correctly recommended spraying in 20 cases, resulting in a sensitivity of 87.0% (95% exact CI: 66.7–97.2%). In all 10 evaluations in which the statistical analysis did not support fungicide application, PedMan correctly recommended no spraying, resulting in a specificity of 100% (95% exact CI: 69.2–100.0%). Accordingly, all 20 spray recommendations were supported by the corresponding statistical analyses, resulting in a PPV of 100% (95% exact CI: 83.2–100.0%). Among the 13 no-spray recommendations, 10 were supported by the statistical analysis, resulting in a NPV of 76.9% (95% exact CI: 46.2–93.8%). The balanced accuracy was 93.5%, the F1 score was 93.0%, and the MCC was 0.818. All three incorrect recommendations were false-negative decisions, in which PedMan recommended withholding fungicide application although a statistically significant difference in leaf abscission was detected between untreated and fungicide-treated trees.

**Table 6.** Contingency table summarizing the validation of PedMan spray recommendations based on field trial and observation outcomes^y^.

|  | Field evaluation<sup>z</sup> |  |  |
|---|---|---|---|
| PedMan recommendation | Spraying warranted | Spraying not warranted | Total |
| Spray | 20 (TP) | 0 (FP) | 20 |
| No spray | 3 (FN) | 10 (TN) | 13 |
| Total | 23 | 10 | 33 |

<sup>Y</sup>Results are from the current (Tables 2 and 4; Figure 3) and the parallel (Gilat et al., 2026) studies
<sup>z</sup>TP = true positive; FP = false positive; FN = false negative; TN = true negative

## 4. Discussion

PedMan was initially developed for olive cultivars highly susceptible to *V. oleaginea* (Ezra et al., 2025; Gilat et al., 2026). In the present study, we extended the system to cultivars differing in susceptibility by adapting the infection-alert criteria according to host resistance. This approach was based on the strategy of Shtienberg et al. (2000), who adjusted rainfall thresholds for chickpea cultivars differing in resistance to *Didymella rabiei*. Because infection by both pathogens is closely associated with rainfall, we evaluated whether the rainfall and temperature criteria established for highly susceptible olive cultivars could be modified according to cultivar susceptibility. Sensitivity analysis indicated that the rainfall threshold of ≥15 mm could be retained across susceptibility classes, whereas the temperature range could be progressively narrowed from 12–18°C for highly susceptible cultivars to 13–17°C for moderately susceptible and 14–16°C for moderately resistant cultivars (Figure 2). Thus, the expansion of PedMan was achieved by maintaining a common rainfall threshold while progressively restricting the temperature conditions under which an infection alert is generated.

The evaluation results support the effectiveness of this approach. PedMan correctly identified all 20 cases in which fungicide application was warranted and correctly recommended no spray in 10 of the 13 cases in which fungicide application was not warranted (Table 6). In highly susceptible cultivars, fungicide application according to PedMan recommendations reduced leaf abscission by 60.8 ± 4.1% (mean ± SE; Gilat et al., 2026), whereas the reduction was 47.9 ± 3.3% in moderately susceptible cultivars (Table 2). In cv. Askal, which was classified as highly resistant, the erroneous no-spray recommendation resulted in a 37.1 ± 3.8% reduction in leaf abscission following fungicide application (Table 4). Had cv. Askal been classified as moderately rather than highly resistant, PedMan would have recommended fungicide application, resulting in a correct recommendation and an effect consistent with those observed for the other two resistance classes. This finding highlights the importance of accurately characterizing cultivar responses to *V. oleaginea* and suggests that refinement of resistance classifications could further improve PedMan performance.

Across all 33 field trials and observations, PedMan correctly classified 30 cases with 90.9% overall accuracy, 87.0% sensitivity, 100% specificity, 100% PPV, 76.9% negative predictive value, 93.5% balanced accuracy, an F1 score of 93.0%, and an MCC of 0.818. The absence of false-positive spray recommendations is particularly important for sustainable disease management, as all 20 spray recommendations were supported by a significant reduction in leaf abscission. Conversely, the three false-negative decisions represent the principal limitation of the system because they indicate missed opportunities for disease control. Given the relatively small validation dataset, the confidence intervals around sensitivity and NPV should be considered when extrapolating these estimates to other cultivars, environments, seasons, and disease-pressure levels.

Despite their potential, the practical application of forecasting models and DSSs for peacock eye disease remains limited, particularly because of the scarcity of independent field evaluation (Thomidis et al., 2021). Consequently, the performance of PedMan could only be compared with prediction models and DSSs developed in other pathosystems. Weather-based models for hop downy mildew achieved 83.3–87.5% sensitivity and 84.4–88.8% specificity (Gent and Ocamb, 2009), while the Nærstad potato late blight model achieved 83% accuracy, 89% sensitivity, and 62% specificity (Hjelkrem et al., 2021). Fusarium head blight models achieved up to 85% accuracy and 100% sensitivity, with 78.4% specificity (Giroux et al., 2016), whereas wheat leaf-blotch DSSs showed substantial interannual variation, with 54–95% correct recommendations depending on year and model (Jørgensen et al., 2020). Compared with these systems, PedMan showed comparable sensitivity but particularly high specificity, indicating a strong ability to avoid unnecessary fungicide applications. This feature is especially relevant because the practical value of a DSS depends not only on predictive accuracy but also on its ability to reduce fungicide use without compromising disease control. Field evaluations of PLANT-Plus and BlightPro demonstrated that DSS-guided programs can reduce or optimize fungicide applications while maintaining disease control and yield (van der Waals et al., 2003; Small et al., 2015), and a meta-analysis of 80 field experiments reported a median 43% reduction in fungicide use without increased disease risk under DSS-based strategies (Lázaro et al., 2021).

The present results therefore place PedMan within the upper range of published plant disease forecasting and DSs, particularly with respect to specificity and the avoidance of unnecessary fungicide applications. Nevertheless, its false-negative decisions indicate that further refinement and evaluation are needed to ensure that the high specificity observed here is maintained without compromising sensitivity. Such evaluation should include a broader range of cultivars, environments, seasons, and disease-pressure levels, with particular attention to cultivar resistance classification and infection-alert criteria.

PedMan has been programmed and made available online, enabling its independent use by Israeli olive growers (https://www.tavazit.com <u>– Hebrew version, on-line translation is evaluable</u>) during the coming growing seasons. Its application may also be relevant to other olive-growing regions characterized by a Csa – hot summer Mediterranean climate (Torres et al., 2017; Fraga et al., 2021). However, before implementation outside Israel, PedMan predictions and spray recommendations should be independently evaluated under local climatic and epidemiological conditions. Such evaluation is particularly important because infection risk depends on the interaction between weather conditions, cultivar susceptibility, and pathogen epidemiology, and because the performance of forecasting systems may vary substantially among environments.

## Acknowledgments

We wish to thank the extension officers Reuven Birger, Mugira Yunis and Nizar Abed-Al-Ahadi who contributed to this research; Nir Kilman and Navot Razon, the growers who implemented PedMan in their olive groves. This research was supported in part by the Israeli Council of Fruit Trees, and the Chief Scientist of Israel Ministry of Agriculture grant #20-02-0039.

## References

Abuamsha, R., Abueid, M., Hajjeh, H., Salman, M., 2013. Evaluation of the incidence and severity of olive leaf spot caused by *Spilocaea oleagina* in different olive cultivars in Palestine. J. Agric. Environ. Int. Dev. 107, 201–212.

Abuley, I.K., Nielsen, B.J., 2019. Integrating cultivar resistance into the TOMCAST model to control early blight of potato, caused by *Alternaria solani*. Crop Prot. 117, 69–76.

Afiad-Lachger, Z., El Guilli, M., Lahlali, R., Amrani, S., 2020. Evaluation of the susceptibility of some Moroccan and international olive cultivars to *Spilocaea oleagina* under field conditions. Plant Pathol. J. 36, 415–424.

Agrios, G.N., 2005. Plant Pathology, fifth ed. Elsevier Academic Press, Burlington, MA, USA.

Altman, D.G., Bland, J.M., 1994. Statistics Notes: Diagnostic tests 1: Sensitivity and specificity. BMJ 308, 1552. 10.1136/bmj.308.6943.1552.

Bernaschina, Y., Alaniz, S., Conde-Innamorato, P., Leoni, C., 2019. Response of olive (*Olea europaea*) cultivars against *Venturia oleaginea* causing olive scab in Uruguay. Agrociencia Uruguay 23, e59. 10.31285/AGRO.23.1.13.

Buonaurio, R., Almadi, L., Famiani, F., Moretti, C., Agosteo, G.E., Schena, L., 2023. Olive leaf spot caused by *Venturia oleaginea*: An updated review. Front. Plant Sci. 13, 1061136. 10.3389/fpls.2022.1061136

Chicco, D., Jurman, G., 2023. The Matthews correlation coefficient (MCC) should replace the ROC AUC as the standard metric for assessing binary classification. BMC Bioinform. 16, 4. 10.1186/s13040-023-00322-4.

Ciccarese, F., Ambrico, A., Longo, O., Schiavone, D., 2002. Search for resistance to *Verticillium*-wilt and leaf spot in olive. Acta Hortic. 586, 717–720.

Estudillo, C., Pérez-Rial, A., Guerrero-Páez, F.A., Díez, C.M., Moral, J., Die, J.V., 2025. Characterization of olive-resistant genes against *Spilocaea oleagina*, the causal agent of scab. Agronomy 15, 452.10.3390/agronomy15020452

Ezra, D., Ygzao, D., Gilat, Y., Shtienberg, D., 2025. Peacock’s eye disease of olive, do we finally understand how to manage it effectively? Phytopathol. Mediterr. 64, 354 (Abstract).

Foody, G.M., 2023. Challenges in the real-world use of classification accuracy metrics: From recall and precision to the Matthews correlation coefficient. PLoS One 18, e0291908. 10.1371/journal.pone.0291908.

Fraga, H., Moriondo, M., Leolini, L., Santos, J.A., 2021. Mediterranean olive orchards under climate change: A review of future impacts and adaptation strategies. Agronomy 11, 56. 10.3390/agronomy11010056.

Gent, D.H., Ocamb, C.M., 2009. Predicting infection risk of hop by *Pseudoperonospora humuli*. Phytopathology 99, 1190–1198. 10.1094/PHYTO-99-10-1190.

Gilat, Y., Ygzao, D., Shtienberg, D., Ezra, D., 2026. Evaluation of the peacock eye disease management (PedMan) system in highly susceptible olive cultivar and using the system for addressing key biological questions. Front. Agron., submitted. 10.64898/2026.06.02.729514

Giroux, M.E., Bourgeois, G., Dion, Y., Rioux, S., Pageau, D., Zoghlami, S., Parent, C., Vachon, E., Vanasse, A. 2016. Evaluation of forecasting models for Fusarium head blight of wheat under growing conditions of Quebec, Canada. Plant Dis. 100, 1192–1201. 10.1094/PDIS-04-15-0404-RE.

Graniti, A., 1993. Olive scab: A review. EPPO Bull. 23, 377–384.

Hjelkrem, A.G.R., Eikemo, H., Le, V.H., Hermansen, A., Nærstad, R. 2021. A process-based model to forecast risk of potato late blight in Norway (the Nærstad model): model development, sensitivity analysis and Bayesian calibration. Ecol. Model. 450, 109565. 10.1016/j.ecolmodel.2021.109565.

Jørgensen, L. N., Matzen, N., Ficke, A., Nielsen, G.C., Jalli, M., Ronis, A., Andersson, B., Djurle, A. 2020. Validation of risk models for control of leaf blotch diseases in wheat in the Nordic and Baltic countries. Eur. J. Plant Pathol. 157, 599–613. 10.1007/s10658-020-02025-6.

Lázaro, E., Makowski, D., Vicent, A. 2021. Decision support systems halve fungicide use compared to calendar-based strategies without increasing disease risk. Comm. Earth Environ. 2, 224. 10.1038/s43247-021-00291-8.

López-Doncel, L.M., García-Berenguer, A., Trapero, A., 1999. Resistance of olive tree cultivars to leaf spot caused by *Spilocaea oleagina*. Acta Hortic. 474, 549–553.

MacDonald, A.J., Walter, M., Trought, M., Frampton, C.M., Burnip, G., 2000. Survey of olive leaf spot in New Zealand. N. Z. Plant Prot. 53, 126–132.

Mahfoud, Y., Choumane, W., Doai, F., 2018. Screening of some olive cultivars (*Olea europaea* L.) for tolerance to Spilocaea oleagina. Int. J. Agric. Environ. Res. 4, 23–30. 10.53555/eijaer.v3i2.25.

Martin, G.C., Sibbett, G.S., 2005. Botany of the olive, in: Sibbett, G.S., Ferguson, L. (Eds.), Olive Production Manual, second ed. University of California, Agriculture and Natural Resources, pp. 15–19.

Moral, J., Alsalimiya, M., Roca, L.F., Díez, C.M., León, L., de la Rosa, R., Barranco, D., Rallo, L., Trapero, A., 2015. Relative susceptibility of new olive cultivars to *Spilocaea oleagina*, *Colletotrichum acutatum*, and *Pseudocercospora cladosporioides*. Plant Dis. 99, 58–64.

Najafi, S., Saremi, H., Jafary, H., Dadras, A., 2023. Evaluation of the relative resistance of different olive cultivars to olive peacock spot disease caused by *Venturia oleaginea*. J. Plant Dis. Prot. 130, 361–369. 10.1007/s41348-022-00697-2.

Niks, R.E., Qi, X., Marcel, T.C., 2015. Quantitative resistance to biotrophic filamentous plant pathogens: Concepts, misconceptions, and mechanisms. Annu. Rev. Phytopathol. 53, 445–470.

Obanor, F.O., Jaspers, M.V., Jones, E.E., Walter, M., 2008. Greenhouse and field evaluation of fungicides for control of olive leaf spot in New Zealand. Crop Prot. 27, 1335–1342.

Obanor, F.O., Walter, M., Jones, E.E., Jaspers, M.V., 2011. Effects of temperature, inoculum concentration, leaf age, and continuous and interrupted wetness on infection of olive plants by *Spilocaea oleagina*. Plant Pathol. 60, 190–199.

Parlevliet, J.E., 1976. Partial resistance of barley to leaf rust *Puccinia hordei* III; The inheritance of the host plant effect on latent period in four cultivars. Euphytica 25, 241–248.

Shtienberg, D., Vintal, H., Brener, S., Retig, B., 2000. Rational management of *Didymella rabiei* in chickpea by integration of genotype resistance and curative application of fungicides. Phytopathology 90, 834–842.

Small, I.M., Joseph, L., Fry, W.E., 2015. Evaluation of the BlightPro decision support system for management of potato late blight using computer simulation and field validation. Phytopathology 105, 1545–1554. 10.1094/PHYTO-05-15-0117-R.

Thomidis, T., Michos, K., Chatzipapadopoulos, F., and Tampaki, A., 2021. Evaluation of two predictive models for forecasting olive leaf spot in northern Greece. Plants 10, 1200.

Torres, M., Pierantozzi, P., Searles, P., Rousseaux, M.C., García-Inza, G., Miserere, A., Bodoira, R., Contreras, C., Maestri, D., 2017. Olive cultivation in the Southern Hemisphere: Flowering, water requirements and oil quality responses to new crop environments. Front. Plant Sci. 8, 1830.

van der Waals, J.E., Denner, F.D.N., van Rij, N., Korsten, L. 2003. Evaluation of PLANT-Plus, a decision support system for control of early blight on potatoes in South Africa. Crop Prot. 22, 821–828. 10.1016/S0261-2194(03)00049-8.

Viruega, J.R., Trapero, A., 1999. Epidemiology of leaf spot of olive tree caused by *Spilocaea oleagina* in southern Spain. Acta Hortic. 474, 531–534.

Viruega, J.R., Roca, L.F., Moral, J., Trapero, A., 2011. Factors affecting infection and development of olive leaf spot caused by *Spilocaea oleagina*. Plant Dis. 95, 1139–1146.

Wu, Y., Small, I.M., Fry, W.E., 2017. Evaluation of the BlightPro decision support system for management of potato late blight using computer simulation and field validation. Phytopathology 107, 1247–1255.

Ygzao, D., Shulhani, R., Shtienberg, D., Ezra, D., 2026a. *Venturia oleaginea*, the causal agent of peacock eye disease in olive, maintains monocyclic and polyetic characteristics in a Mediterranean climate. Plant Dis. 110, 338–346.

Ygzao, D., Shtienberg, D., Ezra, D., 2026b. Optimizing fungicide application timing to control olive Peacock Eye disease: Impact on current and succeeding growing seasons. Plant Dis. 110, 327–337.

Ygzao, D., Gilat, Y., Shtienberg, D., Ezra, D., 2026c. Why peacock eye severity varies with olive tree orientation: Defining criteria to predict infection events. Plant Dis., in press.

